# Imaging Cuprizone-Induced Mitochondrial Dysfunction

**DOI:** 10.1101/2020.12.18.423512

**Authors:** Lucille A. Ray, Gardenia Pacheco, Alexandra Taraboletti, Michael C. Konopka, Leah P. Shriver

## Abstract

Cuprizone is a copper chelator that induces mitochondrial dysfunction in myelin-producing oligodendrocytes and hepatic cells. Inhibition of oxidative phosphorylation has been proposed as a potential mechanism, but the exact relationship between shape changes and metabolic alterations is not well-understood. Here we explore how mitochondrial shape influences oxidative phosphorylation rates by performing simultaneous imaging and respiration measurements within intact cells. We observed that MO3.13 cells exposed to cuprizone undergo an initial increase in respiration followed by mitochondrial dysfunction and genetic dysregulation within 8 hours. Oxygen consumption was measured within 30 minutes of treatment and found to be elevated. This increase was followed by swelling of mitochondria over the first 8 hours, but preceded cell death by 24 hours. A transcriptomic analysis of early changes in cellular gene expression identified alterations within the electron transport chain, stress response pathways, and mitochondrial dynamics compared to control cells. These results suggest that pathological mitochondrial swelling is associated with increased oxygen consumption rates leading to transcriptional changes in respiratory complexes and ultimately mitochondrial failure.

## Introduction

Cuprizone (CPZ) intoxication is a well-studied model of reversible demyelination and mitochondrial dysfunction that is used to probe regenerative strategies for multiple sclerosis (Kashani et al., 2014; Vega-Riquer et al., 2017). Feeding of mice results in characteristic mitochondrial pathology that includes swelling, alterations in mitochondrial metabolic pathways, and oxidative stress (Lindner et al., 2009; Petronilli and Zoratti, 1990). Loss of mature myelinating oligodendrocytes within the brain leads to a characteristic demyelination in corpus callosum, cerebellar peduncles, and hippocampus by 6 weeks (Kipp et al., 2009; Wang et al., 2019). In solution, CPZ can chelate copper and this ability has led to the hypothesis that copper depletion may lead to dysfunction of Complex IV or the mitochondrial superoxide dismutase resulting in respiratory failure and cell death (Acs et al., 2013; Messori et al., 2007). However, CPZ treatment of mice leads to widespread changes in gene expression and metabolism (Han et al., 2020; Taraboletti et al., 2017), suggesting that the mode of action of this toxin is more complex.

A well-characterized pathology in CPZ intoxication is the presence of mitochondrial swelling (megamitochondria) in the liver and brain. The size and shape of mitochondria are regulated by distinct mechanisms (fusion and fission) as well as movement of ions across the inner mitochondrial membrane (Liu et al., 2009; Meyer et al., 2017). Impaired mitochondrial fusion and fission has been implicated in the general progression of neurodegenerative diseases (Knott and Bossy-Wetzel, 2008) and has been identified as one of the factors driving production of reactive oxidative species leading to tissue damage (Ruiz et al., 2020; Ruiz et al., 2018).

Here we utilize a novel method that simultaneously measures mitochondrial shape and oxygen consumption in intact cells. This technique keeps mitochondria in their native environment and prevents perturbations in mitochondrial substrate utilization that can occur with commonly-used protocols for mitochondrial purification (Picard et al., 2011; Wettmarshausen and Perocchi, 2017). We measure the average respiration rate for single cells treated with CPZ by using a fluorescence lifetime porphyrin sensor. In the same cells, mitochondrial shape and size are determined by performing 3-dimensional imaging of mitochondria stained with a MitoTracker™ Red dye. We find that CPZ treatment induces an early increase in respiration associated with mitochondrial swelling that is inconsistent with disruption of oxidative phosphorylation. These early alterations in respiration are then followed by later transcriptional changes in proteins involved in oxidative phosphorylation, lipid metabolism, and signaling. Our results point to a more complicated mechanism of toxicity induced by CPZ, where megamitochondria formation is accompanied by early increased respiration that triggers subsequent transcriptional changes in ETC complexes rather than direct inhibition.

## Results

We have previously used the oligodendroglial cell line, MO3.13 cells as a model system to examine CPZ toxicity as the treatment of these cells with CPZ induces cell death and models many of the metabolic alterations observed in the brains of mice fed this toxin (Chen et al., 2014; Taraboletti et al., 2017). A proposed target of cuprizone is Complex IV, which contains two copper centers which could be inactivated due to loss of copper through chelation (Varhaug et al., 2020). However, other mitochondrial targets such as succinate dehydrogenase have been proposed (Picard et al., 2011) and early studies did not detect disruption in cytochrome proteins in megamitochondria (Wakabayashi et al., 1975). In order to investigate direct inhibition of oxidative phosphorylation, we directly tested whether oxygen consumption was altered by cuprizone treatment in living cells. Respiration rate was measured 30-minutes after cells were treated with 1 mM CPZ or ethanol vehicle. Additionally, we also treated cells with rotenone, a complex I inhibitor as a control for inhibition of oxidative phosphorylation. Cells treated with 1 mM CPZ consumed oxygen at a significantly increased rate of 118 attomoles/cellosec compared to 91.3 attomoles/cell·sec for vehicle-treated cells (single factor ANOVA p = 0.03, Fig 1). The increased respiration rate induced by CPZ was significantly different from rotenone-treated cells which had an oxygen consumption rate of 66.5 attomoles/cell·sec. These results indicate that early mitochondrial pathology induced by CPZ leads to increased respiration rather than inhibition of ETC which would arise from blocking activity of the complexes involved in oxidative phosphorylation and precedes overt swelling of mitochondria.

**Figure 1.**
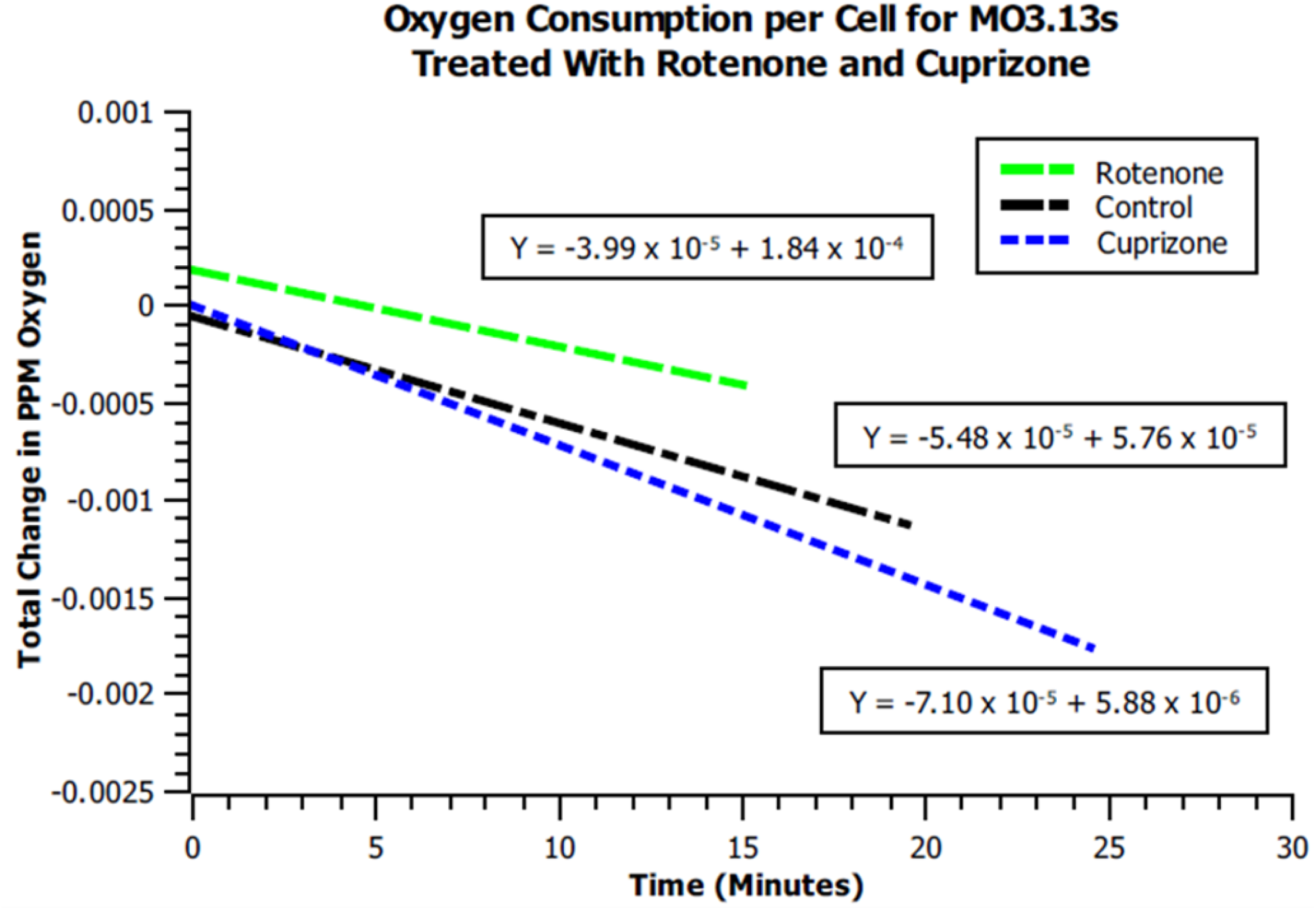
Cuprizone treatment increases respiration in living cells. Oxygen consumption over time for MO3.13 cells treated with 10 μM rotenone, 1 mM cuprizone, and ethanol vehicle. Each slope represents the average oxygen consumption of a single cell. The oxygen consumption rate for ethanol vehicle, 1 mM cuprizone, and 10 uM rotenone was 91.3 attomoles O_2_/cell·sec, 118 attomoles O_2_/cellosec, and 62.4 attomoles O_2_/cell·sec respectively (single factor ANOVA p =.03 for cuprizone versus ethanol vehicle).

We next examined mitochondrial morphology in MO3.13 cells that were pre-treated with 1 mM CPZ or vehicle for 8 hours and then stained with MitoTracker™ Red FM dye for imaging. We chose this time point because we detect transcriptional changes by six hours, but no overt cell deaths. Inverted confocal fluorescence microscopy was used to image cells immediately in Attofluor™ cell chambers. Morphological differences were seen in the mitochondria of CPZ-treated cells when compared to cells treated with vehicle (Fig 2a and b). Mitochondrial crosssectional area was compared for pre-treatment cells and the 8-hour cuprizone treatment cells. CPZ-treated cross-sectional mean area (0.32 μm^2^) was significantly (ANOVA p = 6.7 x 10-6) larger than vehicle-treated cells (0.21 μm^2^, Fig 2c). The enlarged mitochondria were found throughout the cell and many contained visible internal space. Additionally, MitoTracker™ Red FM staining is dependent upon the mitochondrial membrane potential, indicating that these mitochondria are viable and non-depolarized after this treatment period despite swelling.

**Figure 2.**
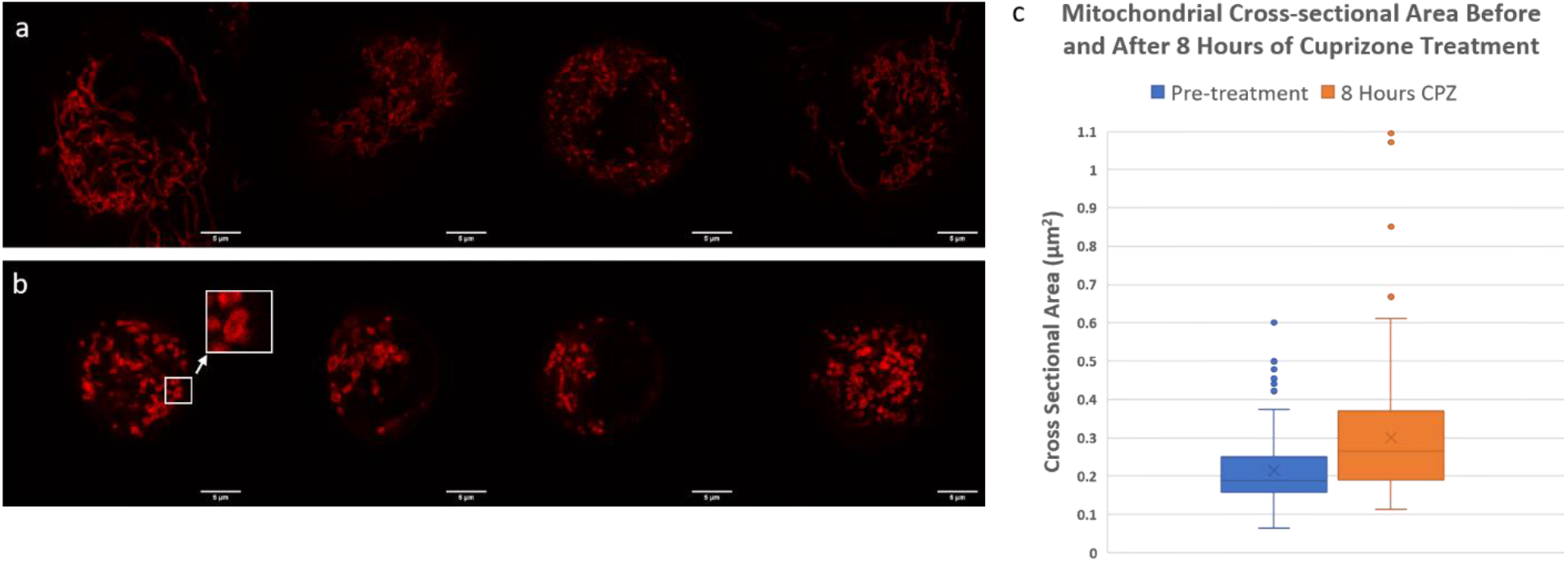
Cuprizone induces mitochondrial swelling prior to cell death: a) Individual MO3.13 cells treated with vehicle (ethanol) for 8 hours then stained with MitoTracker™ Red FM dye. Red staining indicates mitochondria. b) Individual MO3.13 cells treated with cuprizone and ethanol for 8 hours then stained with MitoTracker™ Red FM dye. Red staining indicates mitochondria. Mitochondria with non-stained space within shows swelling (white box). c) Mitochondrial cross-sectional area for pre-cuprizone treated and 8-hour 1 mM cuprizone treated MO3.13 cells. Pre-treatment cross-sectional mean was 0.21 μm^2^ and cuprizone treatment mean was 0.30 μm^2^ (single factor ANOVA p = 6.7 x 10^-6^).

Mice fed CPZ develop focal demyelination in brain regions such as corpus callosum that is accompanied by widespread alterations in brain gene expression (Han et al., 2020). We sought to examine the relationship between the formation of megamitochondria and alterations in cellular gene expression by performing transcriptomics on cuprizone-treated MO3.13 cells compared to vehicle treated cells following six hours of treatment. Overall, CPZ induced 8,054 upregulated and 6,681 downregulated genes (Fig 3a). Pathway analysis of CPZ-treated cells showed that the largest impact of treatment involved cell response to stress, indicating that defects in respiration trigger compensatory pathways to preserve viability. Autophagy, catabolic processes, and apoptosis pathways were also significantly altered after CPZ treatment (Fig 3b). Since SDH and complex IV have been proposed as targets of CPZ, we examined expression of genes associated with the electron transport chain. The proteins that compose all 5 complexes (31 genes downregulated, 6 upregulated), including the adenine nucleotide translocator were significantly downregulated (Fig 3c). Furthermore, complex II was the only complex to be downregulated across all its components. Defects in expression of oxidative phosphorylation genes were also accompanied by downregulation of mitochondrial superoxide dismutase (SOD2), suggesting perturbations in O_2_^•-^ scavenging within mitochondria after CPZ treatment.

**Figure 3.**
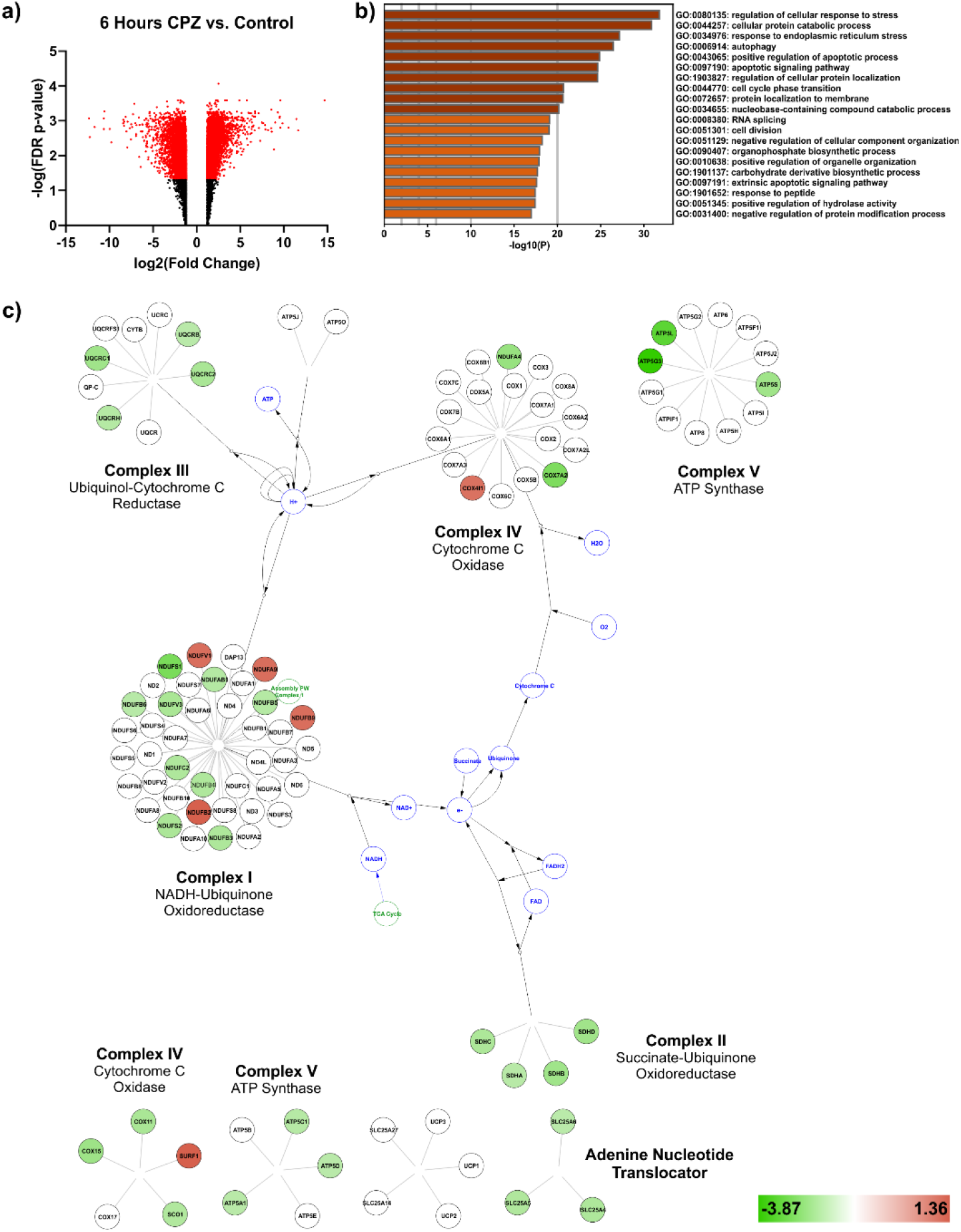
Cuprizone induces transcriptional changes in mitochondrial pathways in MO3.13 cells: a) Volcano plot showing dysregulation of genes in cuprizone-treated cells after 6 hours versus control. Statistically significant dysregulated genes (FDR p < 0.05) are shown in red. b) Top gene ontology (GO) pathways significantly impacted by cuprizone treatment. c) Network clusters of genes upregulated (red) and downregulated (green) in oxidative phosphorylation (ANOVA p < 0.05). For all above, genes with fold changes between −1.2 and +1.2 were not considered.

Imaging of mitochondria in live cells allowed us to investigate the dynamics of the mitochondrial network following the 8-hour CPZ treatment. Mitochondria in MO3.13 cells did not form a consistent network compared to pre-treatment (Fig 4). Instead, they were more sparsely distributed throughout the cell leaving more open space. Transcriptomics confirmed that genes involved in the regulation of the mitochondrial network were perturbed by CPZ treatment. Dynamin-1-like protein (DNM1L) showed downregulation by 2.6 fold while mitochondrial elongation factor 1 (MIEF1) was upregulated and mitofusin 1 and 2 (MFN1, MFN2) were both downregulated. OPA1 and inner membrane mitochondrial protein (IMMT) were both downregulated, both of which drive cristae and inner mitochondrial membrane formation. Increased elongation with decreased fusion may result in an isolated and globular mitochondrial network that we observed by imaging.

**Figure 4.**
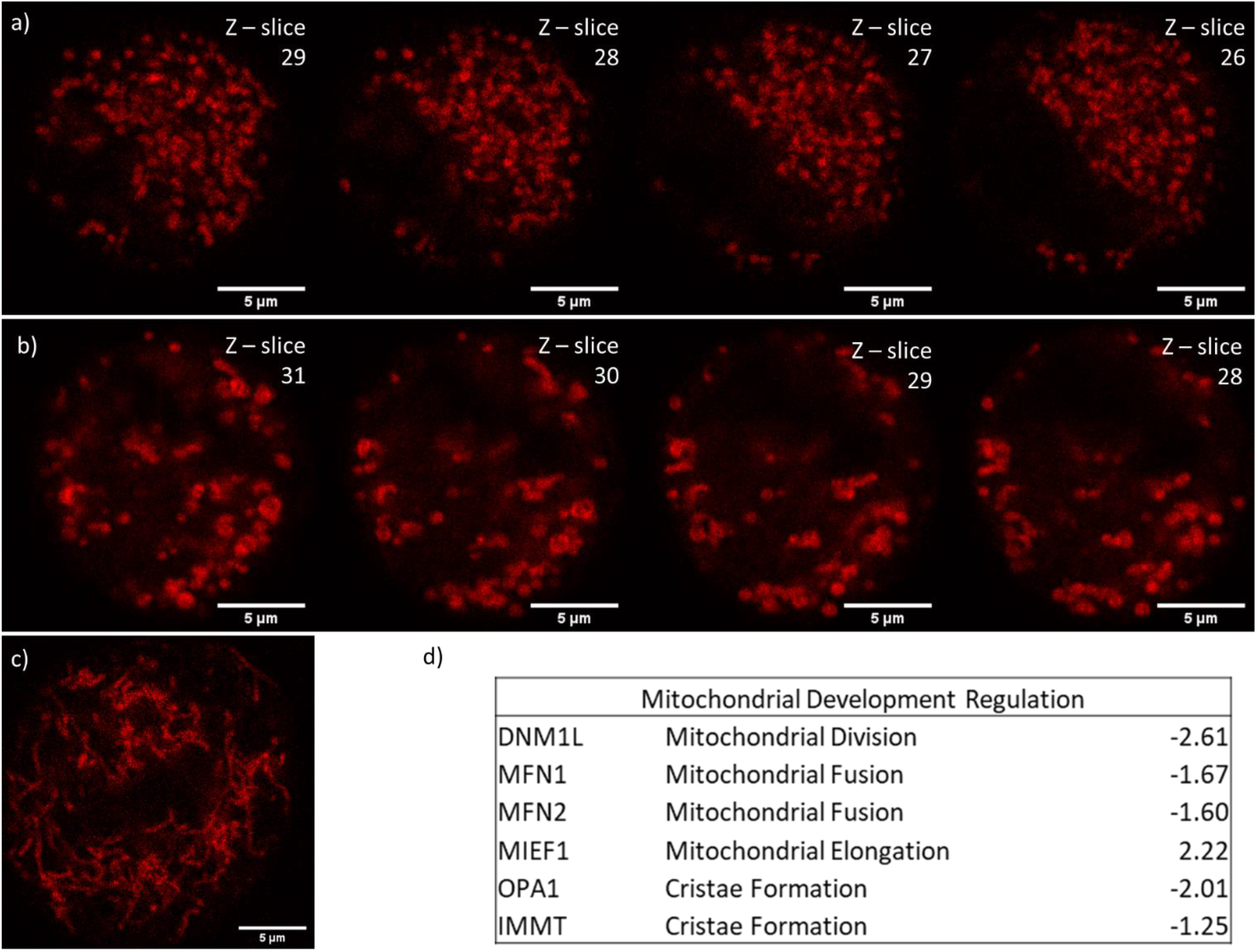
Mitochondrial dynamics are altered by cuprizone toxicity: a) Pre-cuprizone treatment MO3.13 cells stained with MitoTracker™ Red FM dye. Each panel represents a single Z-slice through the cell, moving towards the top of the cell. b) 1 mM cuprizone treated MO3.13 cells stained with MitoTracker™ Red FM dye at 8 hours. Each panel represents a single Z-slice through the cell, moving towards the top of the cell c) MO3.13 cells treated with ethanol vehicle for 24 hours then stained with MitoTracker™ Red FM dye. d) Table of differentially regulated genes related to mitochondrial development with fold-change values.

## Discussion

CPZ intoxication is a well-established model of demyelination that has been used in pre- clinical testing of new regenerative therapies (de Rosa et al., 2019; McMurran et al., 2019; Voskuhl et al., 2019). A complication of this model is that the mechanisms of oligodendrocyte toxicity *in vivo* are multifactorial and potentially involve oxidative stress, energy failure, and inflammation leading to cell death (Vega-Riquer et al., 2017). Here we sought to define early changes in mitochondria that might contribute to oligodendrocyte dysfunction in this model by employing a novel combination of live cell imaging of mitochondrial dynamics coupled to respiration measurements. CPZ treatment initially resulted in an increase in respiration which occurs within one hour of toxin treatment. This response represents an initial response which contrasts with the ability of cuprizone to chelate copper and potentially inhibit respiration (Messori et al., 2007; Taraboletti et al., 2017). Rotenone, by comparison, caused an immediate decrease in respiration consistent with its mechanism of action as a complex I inhibitor. This suggests that CPZ does not completely block oxidative phosphorylation after early administration, but instead may induce a general cellular stress response that increases respiration and the production of reactive oxygen species. This result is consistent with reports of increased oxidative stress *in vivo* (Jakovac et al., 2018; Shiri et al., 2020).

The early increase in respiration following one hour of treatment was subsequently followed by suppression of the expression of oxidative phosphorylation genes and an increase in transcription of genes involved in stress response pathways by six hours of treatment. The reduced activity of complex IV activity in the CPZ mouse model has been attributed primarily to targeted copper chelation as its primary mechanism of action (Acs et al., 2013). Additionally, erythropoientin treatment shows protection against the effects of cuprizone through upregulation of complex IV and a reduction in oxidative stress (Hagemeyer et al., 2012; Kashani et al., 2017). Our *in vitro* findings are consistent with these studies in that we see transcriptional downregulation of the electron transport chain; however, this may be a secondary effect of increased ROS due to stimulation of respiration. In tandem with structural changes in mitochondria, we see expression of proteins involved in regulating mitochondrial fusion and fission. Regulation of DNM1L, MIEF1, MFN1, and MFN2 are typically tightly controlled as part of the cell division process but were all seen to be differentially regulated by CPZ treatment. This suggests that the metabolic disruptions induced by cuprizone may arise from both mitochondrial swelling as well as overall perturbations in the dynamics of the mitochondrial network in cells.

Our results suggest that oligodendrocyte toxicity in cells may arise through a series of events where abnormal respiration leads pathological changes that include mitochondrial swelling and impairment of fusion/fission. This results in activation of multiple stress response pathways in the cell and ultimately downregulation of oxidative phosphorylation and energy failure. While CPZ has the ability to chelate copper in solution, treatment of cells did not result in an immediate inhibition of Complex IV, rather respiratory failure appears to result from perturbations in the expression of the protein complexes involved in oxidative phosphorylation. Finally, imaging mitochondrial dynamics in living cells provides a mechanism for probing metabolic toxins without the need to isolate mitochondria.

## Materials and Methods

### Chemicals

Dulbecco’s Modified Eagle’s Medium (DMEM), Fetal Bovine Serum (FBS), and penicillin streptomycin (P/S) were purchased from Corning (Manassas, VA, USA). HyClone™ DMEM/F-12 (1:1; w/o phenol red) was purchased from GE Healthcare Life Sciences (Logan, UT, USA). Bis(cyclohexanone)oxaldihydrazone (for spectrometric det. of Cu; ≥ 99%) was purchased from Sigma Aldrich (St. Louis, MO, USA). Non-denatured ethanol (200 Proof) was purchased from Decon Laboratories Inc. (King of Prussia, PA, USA). Rotenone (≥ 97%) was purchased from MP Biomedicals (Solon, OH, USA).

### Cuprizone Treatment

MO3.13 oligodendrocyte cell line was cultured in DMEM supplemented with 10% FBS and 1% P/S and maintained in a humified environment at 37 °C with 5% CO2. Unless stated otherwise, complete media contains 10% FBS and 1% P/S. MO3.13 cell solutions were prepared at a final density of 1.0 x 10^4^ cells in complete DMEM/F12 (+2.5% ethanol). 10 mM cuprizone stock solutions were prepared in serum-free DMEM/F12 (+25% ethanol). In the mitochondrial swelling experiments, CPZ solutions were prepared in complete DMEM/F12 (+25% ethanol). These stock solutions were sonicated in a heated FS-14 Solid State/Ultrasonic (Fisher Scientific, USA) water bath until fully dissolved. During this period, the solution was vortexed for 30 sec after 30 min increments. For cell studies, the 10 mM CPZ stock solution was diluted to a final concentration of 1 mM.

### Oxygen Consumption Measurement

MO3.13 cells were incubated on a 3.2 μL volume glass well plate for 15 minutes in serum-free DMEM. Media was removed and replaced with DMEM containing 1 mM cuprizone and 2.5% ethanol. The glass well plate was sealed using a #1.5 thickness 22×22mm coverslip, pre-treated with FluoSpheres carboxylate modified .04 μm Pt luminescent beads on the well-facing side. Oxygen consumption was monitored via the fluorescent lifetime of the beads (ex 390/em 650), collected every 30 seconds. Detection was accomplished using an Andor iStar CCD detector DH743-18-F-A3 with a BNC Model 557 Pulse Generator providing excitation pulses. After oxygen consumption measurement was completed, total cell count was taken via imaging the entire well.

### Mitochondrial Size Measurements

All images were collected on a Nikon Ti-E inverted microscope with the Nikon A1 confocal system using a 100x Plan Apo λ (NA 1.45) oil objective. Excitation was at 561 nm, emission was collected from 570 nm to 620 nm. MO3.13 cells were treated with either 1 mM cuprizone in DMEM containing 2.5% ethanol. Treated cells were stained with MitoTracker^®^ Red FM dye (ThermoFisher) 30 minutes prior to imaging. Individual cells were imaged from top to bottom via 0.5 μm step z-stack. Image data was analyzed to compare mitochondrial size between cells prior-to and after treatment with cuprizone over the course of 8 hours. MO3.13 cells were imaged after exposure to identical media conditions without cuprizone for 24 hours as a control.

### Transcriptomics Analysis

MO3.13 cells were plated at 1.0 x 10^7^ cells in a T25 flask and treated with 1mM cuprizone or vehicle for 6 hours. Three days following treatment, cells were removed and pelleted. Total RNA and microRNA were extracted using a miRNeasy^®^ Mini Kit (Qiagen, Hilden, Germany). Samples were processed by Case Western Reserve Genomics on an Affymetrix^®^ Human Genome U219 microarray. Analysis of the microarray data was performed using the Transcriptome Analysis Console (TAC) software. All genes with p ≤ .016 and q ≤ .050 were considered during analysis (false discovery rate of 5%). Genes were considered to have a positive fold change if the magnitude of change was greater than +1.2-fold, or a negative fold change if the magnitude of change was lower than −1.2-fold. Genes between +1.2-fold and −1.2- fold change were considered no change. Differentially regulated pathways were identified via Metascape.

## Acknowledgements

This work was supported by 1R15 GM119074-01 (L.P.S).

